# Tile-X: A vertex reordering approach for scalable long read assembly [Proceedings]

**DOI:** 10.1101/2025.04.21.649853

**Authors:** Oieswarya Bhowmik, Ananth Kalyanaraman

## Abstract

Traditional approaches for long read assembly compute overlapping reads and subsequently use that overlap information to assemble the contigs. Inherent to this approach is the subproblem of ordering the reads as per their (unknown) genomic positions of origin. However, existing approaches are not designed to explicitly target computation of this true ordering during the assembly process; instead the ordering information becomes available only after the assembly is complete. In this paper, we posit that prior computing of a reliable read ordering, even if imperfect, can significantly reduce the computational burden of the assembly process, preserve assembly quality, and enhance parallel scalability. Specifically, we present `Tile-X`, a novel graph-theoretic vertex reordering-centric approach to compute long read assemblies. The main idea of the approach is to efficiently compute an overlap graph first, use the overlap graph to (re)order the reads (vertices of the graph), and use that ordering to generate a partitioned parallel assembly. We test this idea with two classes of vertex reordering schemes: a) one that uses standard graph vertex reordering schemes that maximize graph locality or bandwidth measures; and b) another class where we custom define a sparsified reordering scheme that exploits sequence characteristics of the underlying graph to reduce the memory and time-footprint for the final assembly step. Using experiments on a combination of real-world and simulated PacBio High Fidelity (HiFi) long reads generated from real genomes, we demonstrate that the `Tile-X` approach is able to achieve substantial improvements over state-of-the-art long read assemblers, in memory efficiency and runtime, while preserving assembly quality metrics such as NGA50 and largest alignment. On average, across all the inputs, `Tile-X` achieved an NGA50 between 1.06*×* and 2.1*×* larger than state-of-the-art assemblers we compared with, while reducing runtime by up to 3.5*×* and memory consumption up to 3.3*×*.

## 1 Introduction

The recent emergence and rapid evolution of long read sequencing technologies has spawned off a new era in the genome discovery, variant discovery, disease identification, and phylogenetic analysis [1, 2, 3, 4, 5]. Single molecule sequencing (SMS) technologies including Oxford Nanopore Technologies (ONT) [2] and Pacific Biosciences (PacBio) [6] are starting to provide long reads covering different length ranges and quality. PacBio provides reads of varying length intervals (30–60 Kbp for CLR, 10–25 Kbp for HiFi) and low error rates (5%–13% for CLR, <1% for HiFi) [3, 5]. ONT offers longer reads (10 Kbp to >100 Kbp) but with higher error rates (8%-15%) [4, 7, 8]. Owing to the rapid evolution of the long read technology, long read assembly remains an actively pursued problem for algorithmic development and optimization.

There are broadly two classes of long read assemblers: those that construct a de Bruijn graph [9, 10] and those that use the overlap layout consensus (OLC) approach [11, 12, 13, 14, 15]—with a majority following the OLC approach. In the OLC approach, overlapping reads are detected (either through alignment-based or alignment-free) methods, and the overlap information is used to build a read graph (or string graph) where nodes are reads and edges are between pairs of overlapping reads. Subsequently, this graph is processed to generate contigs that constitute the output assembly. The individual assemblers typically vary in the way they compute the overlaps and the way they process the graphs to generate contigs. For instance, MECAT [11] leverages a pseudolinear alignment scoring algorithm to accelerate overlap detection, reducing computational overhead substantially. Falcon [12] utilizes a hierarchical genome assembly process to produce phased diploid assemblies that are both accurate and contiguous. Peregrine [13] employs sparse hierarchical minimizers to streamline the overlap detection process, enabling rapid assembly of high-coverage datasets with reduced computational resources. Tools such as `HiCanu` [14] and `Hifiasm` [15] enhance the quality of their alignments through strategies like haplotype phasing and homopolymer compression. However, across all these assemblers the two major phases are the computation of overlap and the use of that overlap information to produce the assemblies.

Inherent to the OLC approach of assembly is the subproblem of ordering the reads as per their (unknown) genomic positions of origin. In fact, if the true ordering of reads along the target genome becomes known, then the problem of *de novo* assembly is significantly simplified because all that remains is to compute the alignment between successive pairs of overlapping reads defined by that order. However, the ordering of reads is not known *a priori*. Furthermore, existing assemblers are not designed to explicitly target computation of this true ordering *during* the assembly process. Instead, the ordering information becomes available after the final assembly is produced. There have been some recent works which have exploited clustering or partitioning ideas to generate an assembly [16]. Clustering tasks can be viewed as an alternative to generate an ordering, binning the reads into buckets, which can be independently processed for assembly. However, generating an ordering (partial or total) has further information that can be exploited.

### Contributions

In this paper, we evaluate the merits of computing an explicit ordering of reads during the early stages of the assembly process. In particular, we posit that computing an explicit read ordering, even if imperfect, has the potential to significantly reduce the computational burden of the assembly process, without compromising on the assemby quality, while enhancing parallel scalability. Specifically, we present `Tile-X`, a novel graph-theoretic vertex reordering-centric approach to compute long read assemblies. The main idea of the approach is to efficiently compute an overlap graph first, use the overlap graph to (re)order the reads (vertices of the graph), and use that ordering to generate a parallel partitioned assembly.

Here, we note that it may not be practical to compute the true ordering. In fact, under various sequence-agnostic graph-theoretic optimization measures (evaluated in [17]), the problem of computing such an optimal ordering is also intractable. However, it is also *not necessary* to compute a perfect ordering. Instead, a cheaper approximate ordering could suffice to generate a coarse partitioning of reads, providing a two-fold benefit: a) separation of reads from unrelated parts of the genome into different partitions (important for reducing potential misassemblies); and b) introducing a way to process the different partitions in parallel, thereby improving scalability.

We test this main idea of using ordering to aid long read assembly, under two classes of vertex reordering schemes: a) one that uses standard graph *vertex reordering* schemes [18, 19, 20] that maximize various graph locality/bandwidth measures [17]; and b) another class where we custom-define a new type of *sparsified reordering* scheme that exploits sequence characteristics of the underlying graph to reduce the computational footprint for the final assembly step. Our experiments on a combination of real-world and simulated PacBio High Fidelity (HiFi) long reads generated from real genomes demonstrate that the `Tile-X` approach is able to achieve substantial improvements over state-of-the-art long read assemblies, in memory efficiency and runtime, while preserving assembly quality metrics such as NGA50 and largest alignment. For instance, for the full human genome, the sparsified reordering version of `Tile-X` (called `Tile-Far`) achieves an NGA50 that is 2.1× larger compared to `Hifiasm`, at a run-time that is 1.9× faster, while consuming only 30% of the memory. Our results also elucidate the inherent performance-quality trade-offs among the ordering schemes. We note that our `Tile-X` approach can be also viewed as a framework because it is generic enough to allow the use of any long read assembler in the last step of contig assembly.

## 2 Methods

In this section, we first outline the major steps of our `Tile-X` workflow for long read assembly using read ordering (Section 2.1). We then provide two problem formulations for the read ordering, and describe our method for each of those formulations (Section 2.2). In Section 2.2.3, we present the design of our overall `Tile-X` parallel implementation.

### Notation

For rest of the paper, we use the following notation. Let *L* = {*r*_1_, *r*_2_, …, *r*_*n*_} denote the set of *n* input long reads. Given a read *r*, the length of the read is denoted by |*r*|. We denote an overlap-based read graph as *G*(*V, E*), where *V* = *L* (i.e., one vertex per read) and an edge (*i, j*) connecting the vertices corresponding to *r*_*i*_ and *r*_*j*_. The graph is undirected. We also associate a numerical weight associated with each edge, denoted by *ω*_*i,j*_, to reflect the strength of overlap between the two reads.

### 2.1 Overview of the Tile-X workflow

As introduced and motivated in Section 1, the main idea of our approach is to compute a read ordering from the read graph (*G*(*V, E*)) and use that ordering information to generate a genome assembly. Below, we present a high-level summary of the major steps (Figure 1), with details presented in subsequent sections: S1) Graph construction: Given *L*, construct a read graph *G*(*V, E*) as per the overlap detection method of choice. Different assemblers choose different methods (alignment-based or alignment-free or a combination). For testing purposes, we use `JEM-mapper` [21, 22]—an alignment-free, distributed, and parallel mapping tool that is both efficient and accurate. `JEM-mapper` employs a sketch-based approach, generating minimizer-based Jaccard sketches to identify overlaps. To further optimize the mapping process, we focus on mapping only the end segments of the long reads—i.e., for each read, we extract a segment of ℓ base pairs (ℓ = 2*Kbp* used in our experiments) from both ends to generate sketches. The output of this step is a read graph *G*(*V, E*) for all the long reads.

**Figure 1.**
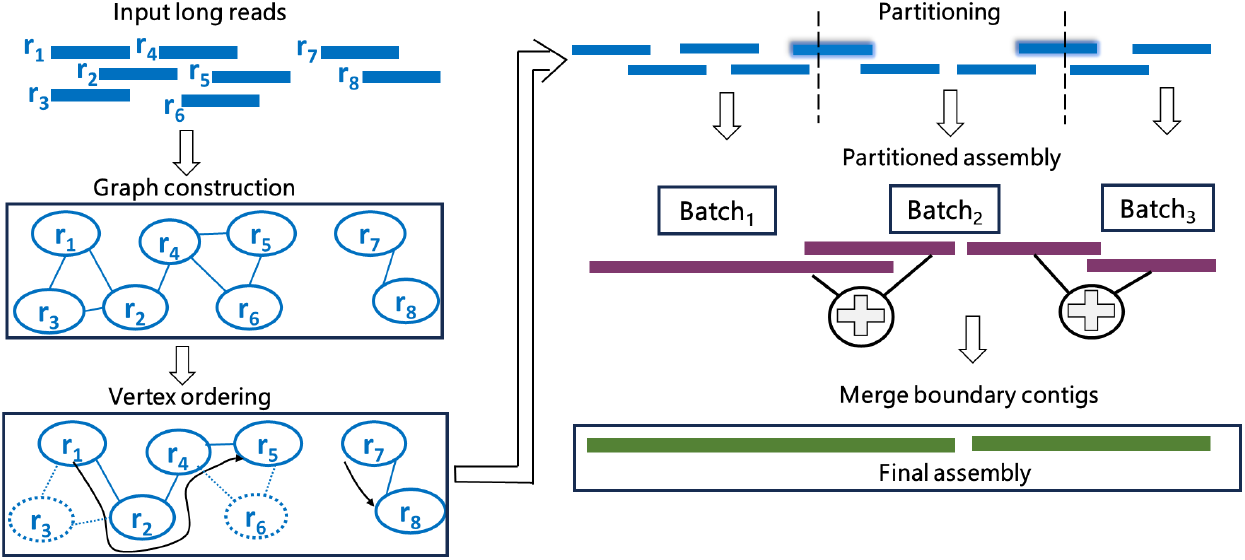
A schematic illustration of the major phases of the proposed `Tile-X` approach.

#### S2) Vertex ordering

From *G*(*V, E*) with *n* vertices (or reads), we generate a vertex ordering *π* : *V*→ {1, 2, …, *n*}, which is a bijective mapping such that the rank of *r*_*i*_ is represented by *π*(*i*). This ordering represents a linear permutation (or a linear ordering) of the input reads. In Section 2.2, we describe various schemes for generating an ordering.

#### S3) Partitioning

Next, using the linear ordering generated in *π*, we partition the read set into *p* subsets. Partitioning can be done in multiple ways: one approach is to identify weak links in the ordering and use them as partition boundaries, while another approach ensures uniform partition sizes to maintain balanced workloads in a parallel setting. In our implementation, we adopt the latter strategy to optimize load balancing. Specifically, we divide the ordered reads into evenly sized partitions, ensuring that each subset contains approximately the same number of reads. After partitioning, we update the subsets to ensure the ending read of one partition say *P*_*r*_, is replicated in its successive partition *P*_*r*+1_ if it exists (shown as the “ghost” vertices straddling the partition boundaries of Figure 1). This is done so as to maintain contiguity across partitions.

#### S4) Partitioned assembly

Next, treating each partition as an individual assembly task, we apply a long read assembler of choice to assemble and produce contigs from each partition. In our implementation and testing, we used `Hifiasm` [15]. Note that this strategy to do a partitioned assembly on each partition generated by step S3, has two advantages: a) Even if the linear ordering detected by the vertex ordering has imperfections, the assembly task will ignore that and treat each partition as just a collection of reads to assemble. This makes the assembly process robust to ordering errors. b) Each partition represents an independent task that can allow all partitions to be assembled in parallel.

#### S5) Merge assemblies

In a final merge step, we combine the individual assemblies produced by the set of successive partitions. This is achieved by running the assembler once again but only on the output contigs that share long reads between two successive partitions.

### 2.2 Read ordering

There are broadly two categories of read ordering schemes (Figure 2) we explore in this paper. In the first approach, we treat the read ordering problem as a graph-theoretic vertex ordering problem. This allows us to explore various standard vertex-ordering schemes (Section 2.2.1). All these schemes produce an ordering for the entire collection of input reads. Next, with a goal to reduce the inbound computational workload for the downstream partitioned assembly step, we present a sparsification-based approach to the ordering problem that can reduce the number of reads selected as part of the final ordering (Section 2.2.2). Assembly workloads that have read sets generated with a high sequencing coverage are better suited to benefit from this approach.

**Figure 2.**
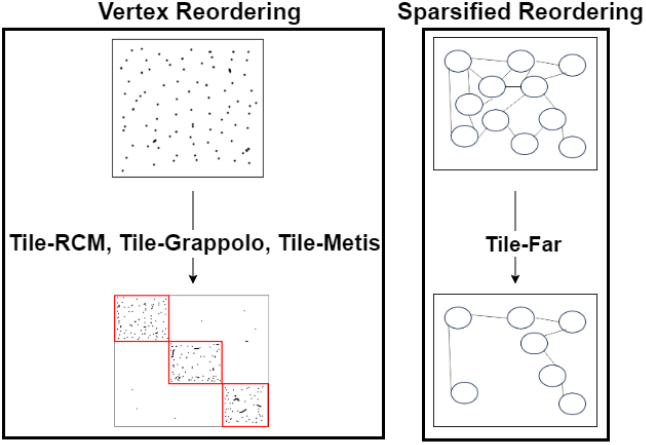
Different vertex ordering schemes of `Tile-X`.

#### 2.2.1 Vertex reordering schemes

Vertex ordering (or reordering) is a classical problem in graph theory and sparse linear algebra used widely to improve locality in computation [17, 23, 24, 25]. Intuitively, given an input sparse matrix, the goal is to reorder the rows such that rows sharing nonzeroes in common appear contiguously, effectively concentrating the nonzeros along the main diagonal of the reordered matrix (as shown in the Figure 2(left)). This reordering formulation also naturally extends to graphs since any graph can be represented as an adjacency matrix (with vertices as rows, and edges as nonzeros). Therefore, reordering is equivalent to renumbering vertices of the graph such that those with shared neighbors in common are contiguously numbered.

More formally, this is the minimum linear arrangement (MINLA) problem [23]: Given a graph *G*(*V, E*), let *π* : *V* → {1, 2, …, |*V* |} be a bijective mapping of verticeLs to a linear permutuation *π* (i.e., ordering). Then the linear arrangement score is given by [26]: *L*(*G, π*) = ∑ _(*i,j*)∈*E*_|*π*(*i*) − *π*(*j*)|. An optimal ordering *π*_*_ is one which minimizes the linear arrangement score for the graph. This optimization problem is NP-Hard [23], and numerous efficient heuristis are used in practice [17].

In the context of genome assembly, it should be easy to see why reordering can be helpful for ordering the vertices of a read graph. Intuitively, because of sequencing coverage, reads that originate from the same region of the genome are likely to share edges to one another in the corresponding read graph. Therefore generating a linear ordering of the vertices of the graph would approximate the ordering of reads along the target genome.

As part of the `Tile-X` framework, we incorporated three vertex ordering schemes that represent three different classes of methods.

- `Tile-RCM`: The Reverse Cuthill-McKee (RCM) [18] ordering scheme is an efficient greedy heuristic that tries to minimize a measure of the graph’s adjacency matrix bandwidth.
- `Tile-Metis`: The Metis ordering is based on a graph partitioner [20], which uses a min-cut multi-level approach to generate a balanced partitioning of vertices (into a pre-specified number of partitions) and subsequently a traversal by each partition to generate its ordering.
- `Tile-Grappolo`: The Grappolo ordering is one that uses the corresponding fast and efficient parallel community detection algorithm [19] to generate an ordering. Intuitively, the first step clusters the vertices into tightly-knit communities using the modularity metric [27], and then traverses the set of vertices by each community to generate an ordering. Under community detection, the number of target communities is determined by the algorithm based on the input (i.e., not user-specified).

While any vertex ordering scheme can be used here, our choices to incorporate and evaluate these three schemes is based on empirical evidence that these schemes outperform other schemes for general graphs [17]. However, their application to read graphs is new in this paper.

#### 2.2.2 Sparsified reordering (Tile-Far)

##### Problem

In this section, we describe a new vertex ordering heuristic called `Tile-Far`. This new scheme is motivated by a goal to reduce the number of reads included in the final ordering. The underlying problem is a variant of the classical ordering problem. Given a graph *G*(*V, E*), the goal is to generate a linear ordering *π*_*s*_ that covers only a “minimal” subset of reads. If *V* ^′^ ⊆*V* denotes that subset, then *π*_*s*_ is a bijective mapping *π*_*s*_ : →{*V* ^′^ 1, 2, …, |*V*|} ^′^. In order to compute *V* ^′^ and a corresponding *π*_*s*_, recall that in the `Tile-X` approach introduced at the start of Section 2.1, the ordering computed in step S2 is subsequently used to generate a partitioned assembly (steps S3 through S5). Therefore, the choice of the subset *V* ^′^ should be such that it is as small as possible (in the interest of performance), while importantly preserving critical overlap information required to generate an accurate assembly. In other words, the goal is to detect a minimum subset *V* ^′^ from which it is possible to generate an assembly that can match the quality of any of the non-sparsified schemes. In order to compute such a minimal subset *V* ^′^ for sparse ordering, we first observe that a read subset *V* ^′^ that corresponds to a small fixed coverage (e.g., 2× or 3 ×) of the target genome should contain sufficient overlap information for assembly. However, the read graph is built using all reads which typically would represent a higher coverage used during sequencing (typically ≥10×). This problem goal is similar to the classical Minimum Tiling Path [28] which was used in the early 2000s to generate a minimal physical map for sequencing experiments.

##### Approach

A potential strategy to generate a sparsified ordering *π*_*s*_ is to follow a two-step process of first sparsifying the input graph (from *G*(*V, E*) to *G*^′^(*V* ^′^, *E*^′^)) and then generating a linear ordering for the *V* ^′^ in *G*^′^. However, this would mean added computational overhead for graph sparsification, which is related to other known NP-Hard problems such as minimum vertex cover [29] and edge cover [30]. Instead of such a two-step process, we present here an efficient algorithm, `Tile-Far`, that directly generates *π*_*s*_ from *G*(*V, E*). Given read graph *G*(*V, E*) and starting from an empty ordering, the `Tile-Far` algorithm incrementally grows its ordering by visiting a subset of vertices in *V* in a certain order and appending them to the current ordering. Initially this traversal starts at a vertex of degree one. In rare instances of a connected component of a graph with no single degree vertices, then the list of all smallest degree vertices in that component are marked as potential “starts” for a traversal. For convenience, we refer to the set of all such start vertices in the entire graph where traversals could start as *V*_S_ (i.e., *V*_𝒮_ ⊆*V*). Let *r*_*i*_ be an arbitrary vertex being visited by the algorithm at any given time. Let *N* (*r*_*i*_) denote the neighbors of *r*_*i*_—i.e., *N* (*r*_*i*_) = {*r*_*j*_ | (*i, j*) ∈ *E*}. From *r*_*i*_, `Tile-Far` tries to greedily find the “farthest” neighbor of *r*_*i*_ from *N* (*r*_*i*_) and append it to *π*_*s*_. More formally, let *r*_*i*_ and *r*_*j*_ be two reads which share a good end-to-end overlap (i.e., semi-global alignment), with the length of their alignment denoted as *Span*(*r*_*i*_, *r*_*j*_).

###### Definition 2.1.

*The* farthest neighbor *of a read r*_*i*_ *is another read r*_*far*_ ∈ *N* (*r*_*i*_) *to which r*_*i*_ *has the maximum span—i*.*e*., *r*_*far*_ = arg max_*j*_ *Span*(*r*_*i*_, *r*_*j*_).

In other words, selecting such a farthest neighbor maximizes the ordering’s span over the target genome in the region corresponding to the two reads. Notably, it has the advantage of skipping over other reads that may lie along the way which also overlap with *r*_*i*_ due to the depth of sequencing coverage (*C*)—thereby generating a sparsification ordering. This idea is illustrated in Figure 3.

**Figure 3.**
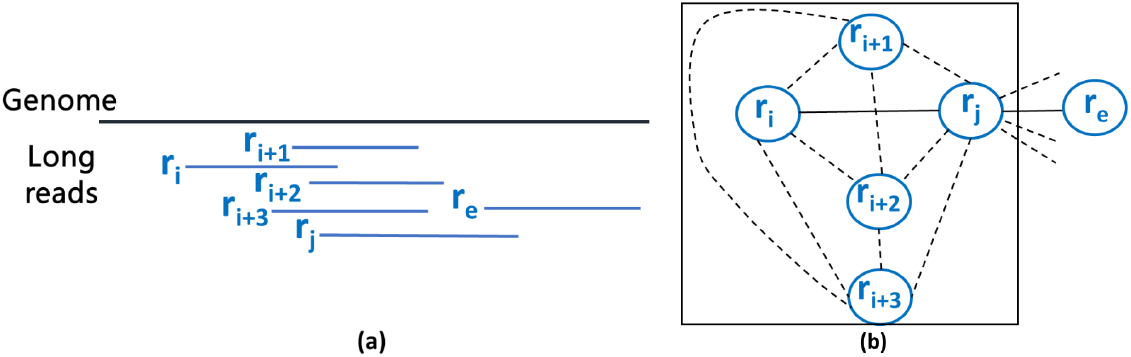
`Tile-Far`: (a) Long reads *r*_*i*_, *r*_*i*+1_, *r*_*i*+2_, *r*_*i*+3_, *r*_*j*_, and *r*_*e*_ spanning overlapping genomic regions; (b) Read graph showing the selection of *r*_*j*_ as the farthest neighbor of *r*_*i*_ (skipping all other reads along the way).

In order to find a farthest neighbor of a given read, without computing alignments and directly from the graph *G*(*V, E*), we start by focusing on maximal cliques within the graph. Let *M* (*r*_*i*_, *r*_*j*_) denote a subgraph of *G*(*V, E*) that is also a maximal clique that contains both *r*_*i*_ and *r*_*j*_. By definition, each read within *M* (*r*_*i*_, *r*_*j*_) shares overlaps with every other read in that clique. The `Tile-Far` heuristic identifies a successor *r*_*j*_ for read *r*_*i*_ by maximizing the following structural gain function:

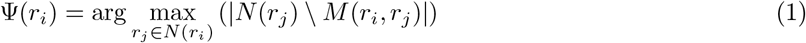

Note that *N* (*r*_*j*_) ⊇ *M* (*r*_*i*_, *r*_*j*_) for any vertex *r*_*j*_. Intuitively, the above structural gain function is a measure of the number of *new* reads that *r*_*i*_ can access through a candidate neighbor *r*_*j*_, that are not already in within its reach in *M* (*r*_*i*_, *r*_*j*_). The more such new reads, the higher the utility of candidate *r*_*j*_ to *r*_*i*_ in extending the contig assembly on the target genome.

While we have posed this approach based on maximal cliques, our algorithm does *not* explicitly compute these cliques as clique computation is hard and expensive in practice [31]. In fact, we can observe from Eqn. 1 we only need to *estimate the cardinality* of the maximal clique size for each (*r*_*i*_, *r*_*j*_) pair in Eqn. 1. Assuming all edges in the input read graph *G*(*V, E*) are “correct”—i.e., true to overlapping reads on the genome—our estimation function is given by |*M* (*r*_*i*_, *r*_*j*_) |= |*N* (*r*_*i*_) ∩ *N* (*r*_*j*_) |. This correctness assumption is not restrictive in practice as argued below. To save runtime further, when our algorithm reaches a vertex *r*_*i*_, our algorithm lazily calculates this intersection cardinality for each of *r*_*i*_’s neighbors at that time, and applies the arg max operation to obtain *ψ*(*r*_*i*_). Algorithm 1 summarizes the main steps of our `Tile-Far` algorithm.

As noted above, the structural gain function objective assumes that the edges in the read graph are “true”, i.e., the reads connected via an edge are truly overlapping on the target genome. However, this depends on the precision accuracy of the mapping procedure (step S1), and there is always a risk of placing a chimeric edge between two reads (belonging to two different parts of the genome). To mitigate the risk of creating a chimeric merge as part of the sparsified ordering, we introduce an additional condition to check before identifying the farthest neighbor *ψ*(*r*_*i*_). If the maximum choice neighbor has a |*M* (*r*_*i*_, *r*_*j*_) | below a certain threshold *τ*, then we instead select the next best choice of neighbor (if one exists) that satisfies this threshold *τ*. This reduces the chance of exposing an unreliable farthest neighbor and alleviates the risk of creating false merges in the subsequent assembly step. If the read sequencing coverage is *C*, then we set 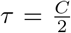 in our experiments.

###### Algorithm 1

`Tile-Far` Heuristic

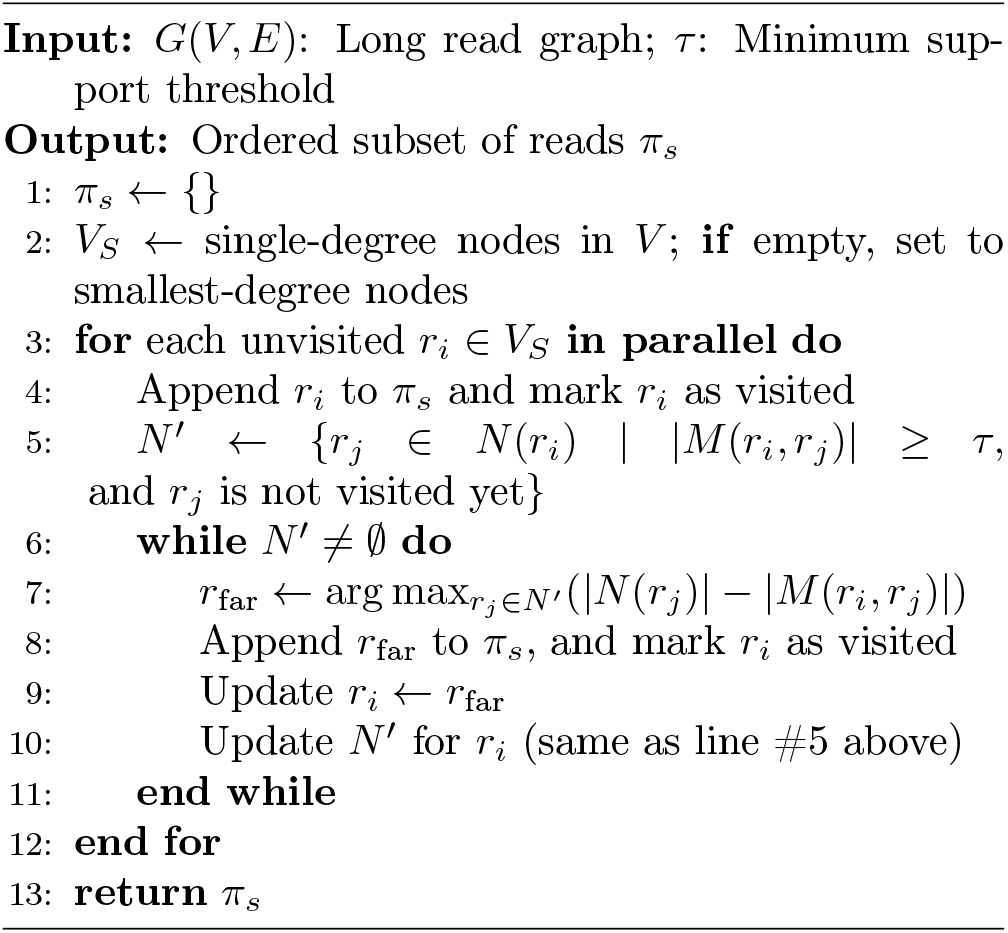

##### Algorithmic properties

Even though the `Tile-Far` algorithm is a heuristic, there are several provable properties of the algorithm. Collectively these properties serve to highlight the algorithm’s advantages and limitations.

###### Lemma 2.1.

*The farthest neighbor relationship ψ*(.) *(as defined in Eqn. 1) is* not *symmetric*.

*Proof*. Consider two reads *r*_*i*_ and *r*_*j*_ which share an edge in *G*(*V, E*). The choice for the farthest neighbor in Eqn. 1 depends on the degree of the destination vertex, which may be different for the two vertices. Therefore, there is no guarantee that if *r*_*j*_ = *ψ*(*r*_*i*_) then *r*_*i*_ = *ψ*(*r*_*j*_).

###### Lemma 2.2.

*Each read will feature in exactly one of the orderings reported by `Tile-Far`*.

*Proof*. The lemma holds because of the `visit` flag maintained at each vertex by the algorithm.

The implications of the above two lemmas is that the `Tile-Far` heuristic is non-deterministic and can potentially generate different output orderings from the same input.

Next we show an important property, about the degree of sparsity that can be expected of `Tile-Far`.

###### Definition 2.2.

*An ordering of m reads π*_*t*_ = [*r*_1_, *r*_2_, … *r*_*m*_] *is referred to as a true ordering if the reads cover a contiguous stretch of the target genome, with every successive pair of reads in the ordering truly overlapping*.

###### Lemma 2.3.

*Consider a true ordering of reads π*_*t*_ : [**r**_**i**_, *r*_*i*+1_, *r*_*i*+2_, …, *r*_*i*+*k*_, **r**_**j**_, *r*_*e*_] *such that r*_*i*_ *overlaps with r*_*j*_ *but there exists no overlap between reads* (*r*_*i*_, *r*_*e*_). *Then, if the Tile-Far algorithm visits r*_*i*_ *then it is likely to identify r*_*j*_ *as its farthest neighbor*.

*Proof*. Given that *π*_*t*_ is a true ordering and given that *r*_*i*_ and *r*_*j*_ share an overlap, we can expect that all the intermediate reads *r*_*i*+1_ through *r*_*i*+*k*_ also share overlaps with one another and with *r*_*i*_ and *r*_*j*_—thereby forming a clique in the read graph *G*(*V, E*) (as illustrated in Fig. 3). Furthermore, since read *r*_*i*_ does not overlap with *r*_*e*_, the collection {*r*_*i*_, *r*_*i*+1_, *r*_*i*+2_, …, *r*_*i*+*k*_, *r*_*j*_} form a maximal clique for the vertex pair (*r*_*i*_, *r*_*j*_), with size *M* (*r*_*i*_, *r*_*j*_) = *k* + 2. This also implies that *M* (*r*_*i*_, *r*_*j*_) = *M* (*r*_*i*_, *r*_*f*_) for *i* + 1≤ *f* ≤*i* + *k*. Therefore, when the `Tile-Far` algorithm considers which of its neighbors from *r*_*i*+1_ through *r*_*j*_ can be considered farthest, the choice is determined by which of those candidate reads *r*_*c*_ maximizes |*N* (*c*) \*M* (*r*_*i*_, *r*_*c*_) | (by Eqn. 1)—which is same as the read that maximizes |*N* (*r*_*c*_) | (since all |*M* (*r*_*i*_, *r*_*c*_) | are the same relative to *r*_*i*_). We observe here that |*N* (*r*_*j*_) | ≥|*N* (*r*_*c*_) |, over all candidates *c*. By contradiction, if there were to be an intermediate read *r*_*f*_, where *i* + 1≤ *f* ≤*i* + *k*, that has a larger vertex degree that would imply that there has to be at least one additional read (such as *r*_*e*_) to the right of *r*_*f*_ that it overlaps with. But if that is the case, then *r*_*j*_, which is further right of *r*_*f*_ in the true ordering would also have to overlap with *r*_*e*_ thereby ensuring that |*N* (*r*_*j*_)| ≥ |*N* (*r*_*c*_)|. Therefore, the `Tile-Far` algorithm would select *r*_*j*_ as *ψ*(*r*_*i*_).

The implication of Lemma 2.3 is that the algorithm is likely to skip all the intermediate reads [*r*_*i*+1_, *r*_*i*+2_, …, *r*_*i*+*k*_], which corresponds to a significant saving in ordering retention as *k* increases.

#### 2.2.3 Parallel implementation

We have implemented the `Tile-X` framework including `Tile-RCM`, `Tile-Grappolo`, `Tile-Metis`, and `Tile-Far` in C/C++ and using the MPI message passing library for communication under the distributed memory model, and OpenMP for shared memory multithreaded parallelism. Owing to space limitations, we skip the inner details of the parallel implementation. Our code is available as open source at https://github.com/Oieswarya/Tile-X.

## 3 Results and Discussion

### 3.1 Experimental setup

#### Test inputs

For our experiments, we used genome inputs downloaded from the NCBI GenBank [32], as summarized in Table 1. We used the Sim-it PacBio HiFi simulator [33], with a default 10× coverage and read length median of 10Kbp. In addition to simulated reads, we also used two real-world HiFi sequencing datasets, both generated using the PacBio Sequel II system: a caddisfly genome (*H. magnus*) [34], and a drumfish genome (*N. coibor*) [35]. **Test platform:** All experiments were conducted on a distributed memory cluster with 9 compute nodes, each with 64 AMD Opteron™ (2.3GHz) cores and 128 GB DRAM. The nodes are interconnected using 10Gbps Ethernet and share 190TB of ZFS storage. The distributed executions of `Tile-X` were performed using MPI with up to *p* = 64 processes (4 compute nodes, each running 16 processes, with 1 thread per process), while the multithreaded executions used 64 threads on a single node of the cluster. For all other the state-of-the-art assemblers, we ran them in their multithreaded mode on 64 threads (mapped to 64 cores) on a single node of the cluster. For all runs, we ran `Tile-X` with a batch size of 16,384 in the batch assembly step.

**Table 1:** Input datasets used in our experiments. All genome inputs were downloaded from NCBI GenBank [32].

| Input genome | | $\mathcal{L}$: Long read statistics (HiFi reads) | | |
| --- | --- | --- | --- | --- |
| Genome | Genome length (bp) | No. long reads (m) | Total length (bp) | Read length (avg. ± std. dev) |
| *D. busckii* | 118,492,362 | 123,781 | 1,258,903,798 | 10,170 ± 3,406 |
| *C. septempunctata* | 398,868,586 | 390,797 | 3,981,681,897 | 10,188 ± 3,398 |
| *B. splendens* | 441,388,503 | 432,230 | 4,404,143,269 | 10,189 ± 3,396 |
| *H. aestivaria* | 501,713,186 | 491,533 | 5,005,317,758 | 10,183 ± 3,397 |
| *P. sinus* | 2,371,540,524 | 2,323,801 | 23,677,646,469 | 10,189 ± 3,398 |
| *H. sapiens* | 3,298,430,637 | 3,083,048 | 31,292,504,943 | 10,149 ± 3,434 |
| *N. coibor* | (620 Mbp est.) | 1,919,461 | 29,259,679,568 | 15,243 ± 2,736 |
| *H. magnus* | (1.2 Gbp est.) | 2,436,589 | 28,013,062,204 | 11,496 ± 720 |

### 3.2 Qualitative evaluation

We compare the `Tile-X` methods against other state-of-the-art assemblers including `Hifiasm` [15], `HiCanu` [14], `HiFlye` [36], and `GoldRush` [10]. All the quality results for are shown in Table 2. Our results show that for all inputs `Tile-X` methods produce comparable or better assembly contiguity (NGA50, NG50, largest alignment) compared to standard tools. This is achieved while maintaining near perfect genome coverage and duplication ratio, and reduced misassemblies (effect of ordering and partitioning in `Tile-X`).

**Table 2:**
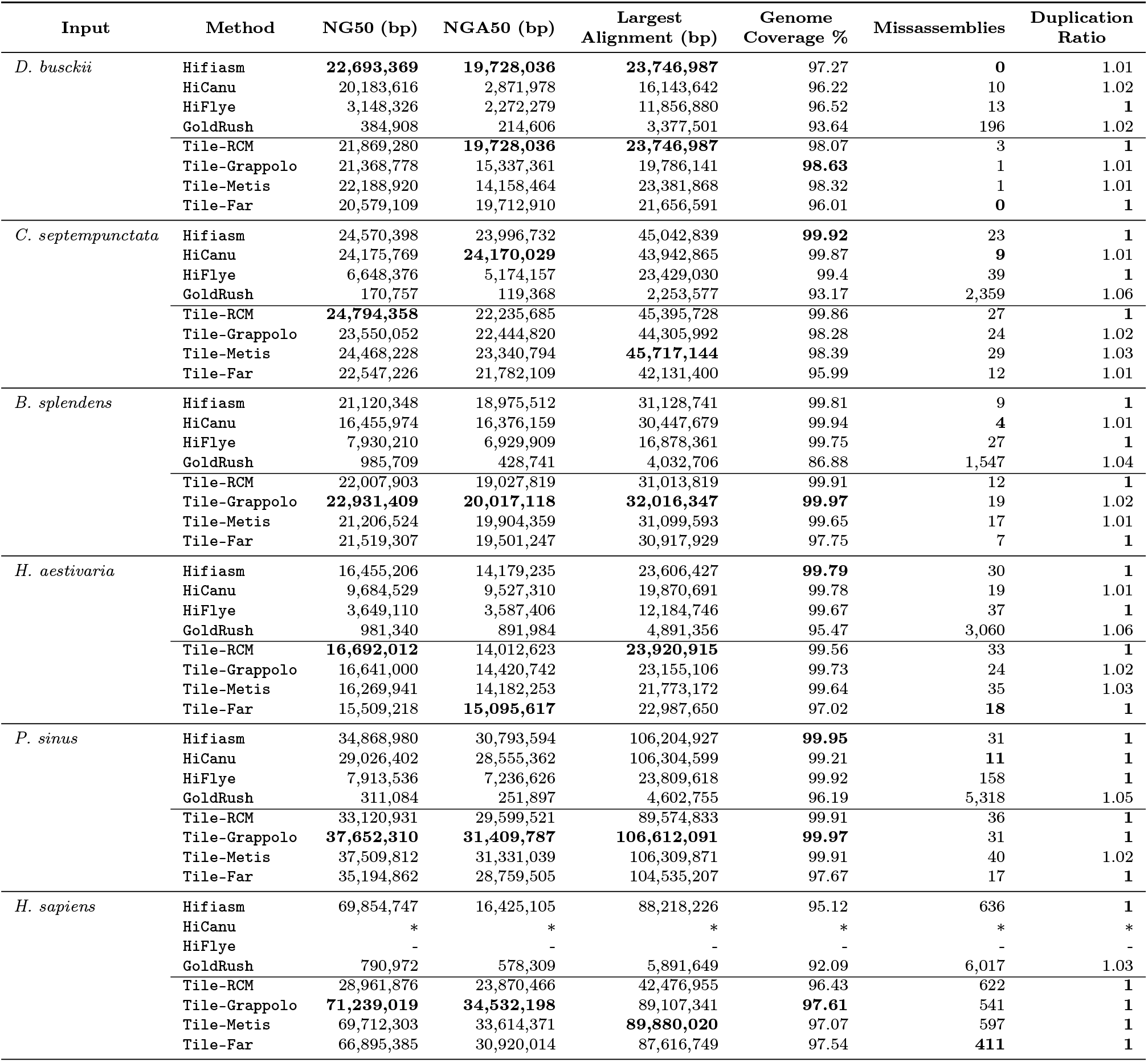
Qualitative comparison of the output contigs generated by the different tools on the different inputs. Symbol * indicates that the corresponding runs did not complete within 6 hours; symbol indicates − that the corresponding runs required more than 256 GB of memory which was the system maximum memory. Bold face values show the best results for each input.

| Input | Method | NG50 (bp) | NGA50 (bp) | Largest Alignment (bp) | Genome Coverage % | Missassemblies | Duplication Ratio |
| --- | --- | --- | --- | --- | --- | --- | --- |
| *D. busckii* | Hifiasm | **22,693,369** | **19,728,036** | **23,746,987** | 97.27 | **0** | 1.01 |
|  | HiCanu | 20,183,616 | 2,871,978 | 16,143,642 | 96.22 | 10 | 1.02 |
|  | HiFlye | 3,148,326 | 2,272,279 | 11,856,880 | 96.52 | 13 | **1** |
|  | GoldRush | 384,908 | 214,606 | 3,377,501 | 93.64 | 196 | 1.02 |
|  | Tile-RCM | 21,869,280 | **19,728,036** | **23,746,987** | 98.07 | 3 | **1** |
|  | Tile-Grappolo | 21,368,778 | 15,337,361 | 19,786,141 | **98.63** | 1 | 1.01 |
|  | Tile-Metis | 22,188,920 | 14,158,464 | 23,381,868 | 98.32 | 1 | 1.01 |
|  | Tile-Far | 20,579,109 | 19,712,910 | 21,656,591 | 96.01 | **0** | **1** |
| *C. septempunctata* | Hifiasm | 24,570,398 | 23,996,732 | 45,042,839 | **99.92** | 23 | **1** |
|  | HiCanu | 24,175,769 | **24,170,029** | 43,942,865 | 99.87 | **9** | 1.01 |
|  | HiFlye | 6,648,376 | 5,174,157 | 23,429,030 | 99.4 | 39 | **1** |
|  | GoldRush | 170,757 | 119,368 | 2,253,577 | 93.17 | 2,359 | 1.06 |
|  | Tile-RCM | **24,794,358** | 22,235,685 | 45,395,728 | 99.86 | 27 | **1** |
|  | Tile-Grappolo | 23,550,052 | 22,444,820 | 44,305,992 | 98.28 | 24 | 1.02 |
|  | Tile-Metis | 24,468,228 | 23,340,794 | **45,717,144** | 98.39 | 29 | 1.03 |
|  | Tile-Far | 22,547,226 | 21,782,109 | 42,131,400 | 95.99 | 12 | 1.01 |
| *B. splendens* | Hifiasm | 21,120,348 | 18,975,512 | 31,128,741 | 99.81 | 9 | **1** |
|  | HiCanu | 16,455,974 | 16,376,159 | 30,447,679 | 99.94 | **4** | 1.01 |
|  | HiFlye | 7,930,210 | 6,929,909 | 16,878,361 | 99.75 | 27 | **1** |
|  | GoldRush | 985,709 | 428,741 | 4,032,706 | 86.88 | 1,547 | 1.04 |
|  | Tile-RCM | 22,007,903 | 19,027,819 | 31,013,819 | 99.91 | 12 | **1** |
|  | Tile-Grappolo | **22,931,409** | **20,017,118** | **32,016,347** | **99.97** | 19 | 1.02 |
|  | Tile-Metis | 21,206,524 | 19,904,359 | 31,099,593 | 99.65 | 17 | 1.01 |
|  | Tile-Far | 21,519,307 | 19,501,247 | 30,917,929 | 97.75 | 7 | **1** |
| *H. aestivaria* | Hifiasm | 16,455,206 | 14,179,235 | 23,606,427 | **99.79** | 30 | **1** |
|  | HiCanu | 9,684,529 | 9,527,310 | 19,870,691 | 99.78 | 19 | 1.01 |
|  | HiFlye | 3,649,110 | 3,587,406 | 12,184,746 | 99.67 | 37 | **1** |
|  | GoldRush | 981,340 | 891,984 | 4,891,356 | 95.47 | 3,060 | 1.06 |
|  | Tile-RCM | **16,692,012** | 14,012,623 | **23,920,915** | 99.56 | 33 | **1** |
|  | Tile-Grappolo | 16,641,000 | 14,420,742 | 23,155,106 | 99.73 | 24 | 1.02 |
|  | Tile-Metis | 16,269,941 | 14,182,253 | 21,773,172 | 99.64 | 35 | 1.03 |
|  | Tile-Far | 15,509,218 | **15,095,617** | 22,987,650 | 97.02 | **18** | **1** |
| *P. sinus* | Hifiasm | 34,868,980 | 30,793,594 | 106,204,927 | **99.95** | 31 | **1** |
|  | HiCanu | 29,026,402 | 28,555,362 | 106,304,599 | 99.21 | **11** | **1** |
|  | HiFlye | 7,913,536 | 7,236,626 | 23,809,618 | 99.92 | 158 | **1** |
|  | GoldRush | 311,084 | 251,897 | 4,602,755 | 96.19 | 5,318 | 1.05 |
|  | Tile-RCM | 33,120,931 | 29,599,521 | 89,574,833 | 99.91 | 36 | **1** |
|  | Tile-Grappolo | **37,652,310** | **31,409,787** | **106,612,091** | **99.97** | 31 | **1** |
|  | Tile-Metis | 37,509,812 | 31,331,039 | 106,309,871 | 99.91 | 40 | 1.02 |
|  | Tile-Far | 35,194,862 | 28,759,505 | 104,535,207 | 97.67 | 17 | **1** |
| *H. sapiens* | Hifiasm | 69,854,747 | 16,425,105 | 88,218,226 | 95.12 | 636 | **1** |
|  | HiCanu | * | * | * | * | * | * |
|  | HiFlye | - | - | - | - | - | - |
|  | GoldRush | 790,972 | 578,309 | 5,891,649 | 92.09 | 6,017 | 1.03 |
|  | Tile-RCM | 28,961,876 | 23,870,466 | 42,476,955 | 96.43 | 622 | **1** |
|  | Tile-Grappolo | **71,239,019** | **34,532,198** | 89,107,341 | **97.61** | 541 | **1** |
|  | Tile-Metis | 69,712,303 | 33,614,371 | **89,880,020** | 97.07 | 597 | **1** |
|  | Tile-Far | 66,895,385 | 30,920,014 | 87,616,749 | 97.54 | **411** | **1** |

As the genome size increases, `Tile-Grappolo` generally outperformed the other methods across most of the metrics. For instance, for *H. sapiens*, `Tile-Grappolo` achieved an NGA50 of 34.5Mbp which is 2.1 × longer than `Hifiasm`. The better results observed for `Tile-Grappolo` (relative to other vertex ordering schemes) is consistent with the relative performance reported on generic graphs in [17]. Largely this is owing to the rigorous optimization function of modularity that it internally uses, without bounding the sizes or number of communities prior to generation of the ordering. However, it is notable that `Tile-Far`, despite sparsifying on the read space, still produced quality comparable to the other `Tile-X` methods.

In general, despite the minor deviations in quality, all `Tile-X` schemes performed comparably, and in many cases also outperformed the state-of-the-art assemblers. Among the existing assemblers, `Hifiasm` in general produced the best outputs. However, in almost all cases (except *D. busckii*), the `Tile-X` implementations produce better quality compared to `Hifiasm`, demonstrating the positive effect of ordering prior to using a standalone assembler. The `Tile-X` methods also produced less misassemblies in multiple cases, demonstrating the value of partitioning after ordering, which would reduce ambiguity in the subsequent partitioned assembly step.

### 3.3 Performance evaluation

Table 3 shows the parallel performance for three of the largest inputs comparing all the tools (results for all inputs is provided in the supplementary materials in Table S1). We see that `Tile-X` consistently demonstrates significant speedups over the state-of-the-art assemblers, and among the `Tile-X` tools, `Tile-Far` is the fastest, demonstrating the value of sparsification even for a small coverage inputs (10×). For instance for *H. sapiens*, `Tile-Far` is 1.9× faster than `Hifiasm`.

**Table 3:**
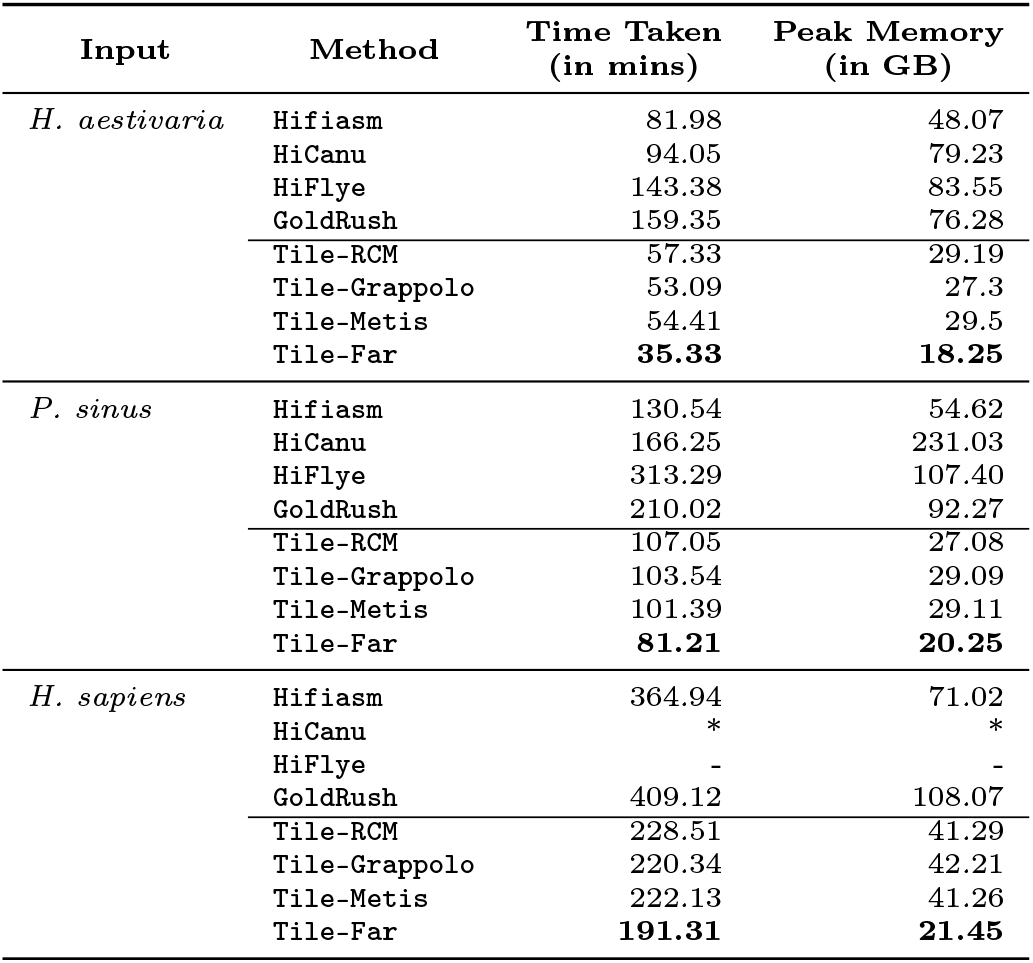
Performance comparison of the output contigs generated by the different tools on the three larger inputs (simulated). Symbol *indicates that the corresponding runs did not complete within 6 hours; symbol − indicates that the corresponding runs required more than 252GB of memory which was the system maximum memory. Bold face values show the best results for each input.

On average `Tile-Far` was 1.9× to 3.5× faster than fastest state-of-the-art tool for any input. The factor improvements with `Tile-Far` correlate to the sparsification factors achieved by `Tile-Far` (data not shown due to space).

Table 3 also shows the memory consumption of all the tools. Here again, `Tile-X` outperforms the other tools, demonstrating the value of breaking down the input through ordering into partitions. Furthermore, among the `Tile-X` schemes, `Tile-Far` consumes the least memory, i.e., between 2.6× and 3.3× less than `Hifiasm`—demonstrating the value of sparsification. We note that both the runtime and memory performance of the `Tile-X` implementations can further improve as we increase the number of processors (due to their distributed memory implementation).

#### Impact of sparsification on high coverage inputs

We note here that the true space saving impact of the sparsification idea in `Tile-Far` can be realized when we start increasing the sequencing coverage. To demonstrate this point, we ran `Tile-Far` over inputs obtained using increasing coverage, ranging from 4× to 30× on the *H. aestivaria* input. Results are shown in Table 4. The results show that the qualitative gains plateau out after 10× coverage. However, neither the runtime nor the memory increases with `Tile-Far` beyond the 9× coverage setting. In fact, with increasing coverage, the sparsification rate only improves (i.e., fraction of vertices retained decreases). These results show the effectiveness of sparsification to reduce redundancy as shown in the property of Lemma 2.3.

**Table 4:**
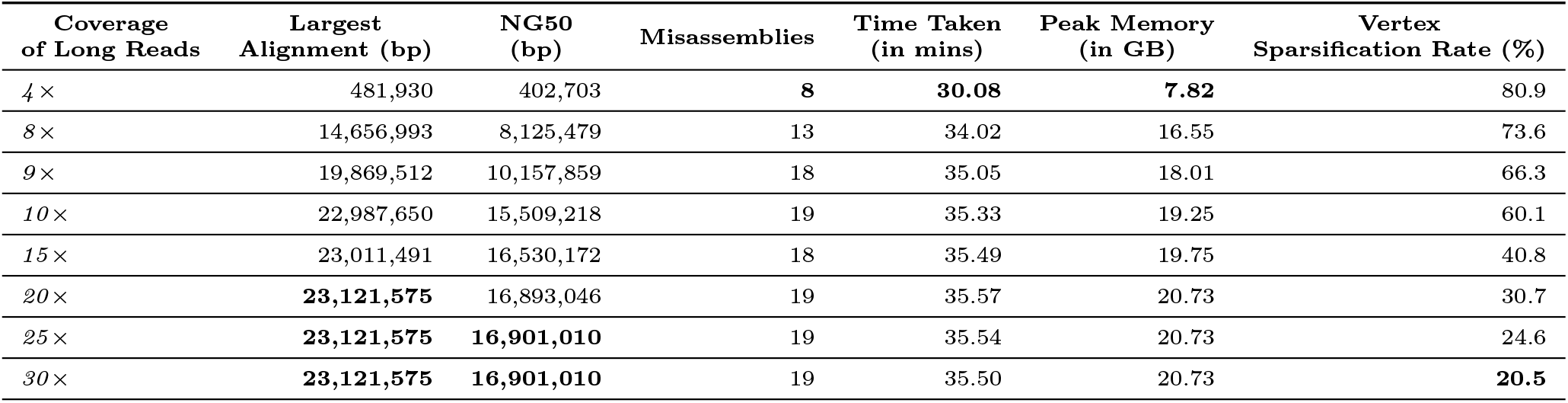
Quality and performance evaluation of running Tile-Far with different coverages of long reads on input *H. aestivaria*. Vertex sparsification rate is measured as the percentage of reads retained in the sparsified ordering relative to the total input. Bold face values show the best results for each metric.

| Coverage of Long Reads | Largest Alignment (bp) | NG50 (bp) | Misassemblies | Time Taken (in mins) | Peak Memory (in GB) | Vertex Sparsification Rate (%) |
| --- | --- | --- | --- | --- | --- | --- |
| 4× | 481,930 | 402,703 | **8** | **30.08** | **7.82** | 80.9 |
| 8× | 14,656,993 | 8,125,479 | 13 | 34.02 | 16.55 | 73.6 |
| 9× | 19,869,512 | 10,157,859 | 18 | 35.05 | 18.01 | 66.3 |
| 10× | 22,987,650 | 15,509,218 | 19 | 35.33 | 19.25 | 60.1 |
| 15× | 23,011,491 | 16,530,172 | 18 | 35.49 | 19.75 | 40.8 |
| 20× | **23,121,575** | 16,893,046 | 19 | 35.57 | 20.73 | 30.7 |
| 25× | **23,121,575** | **16,901,010** | 19 | 35.54 | 20.73 | 24.6 |
| 30× | **23,121,575** | **16,901,010** | 19 | 35.50 | 20.73 | **20.5** |

### 3.4 Real-world dataset evaluation

We evaluated the performance of `Tile-X` on two real-world HiFi sequencing datasets as shown in Table 5. In terms of assembly quality, `Tile-X` showed significant improvements over state-of-the-art assemblers. For the *H. magnus* dataset, `Tile-RCM` achieved a 1.2× improvement in N50 over `Hifiasm`, and a 18.1× improvement over `HiFlye`. For the input *N. coibor*, `Tile-Grappolo` outperformed `Hifiasm` by 1.3× in N50. These results highlight the quality improvements that `Tile-X` provides in real-world applications, demonstrating its potential as a powerful tool for large-scale genome assembly.

**Table 5:**
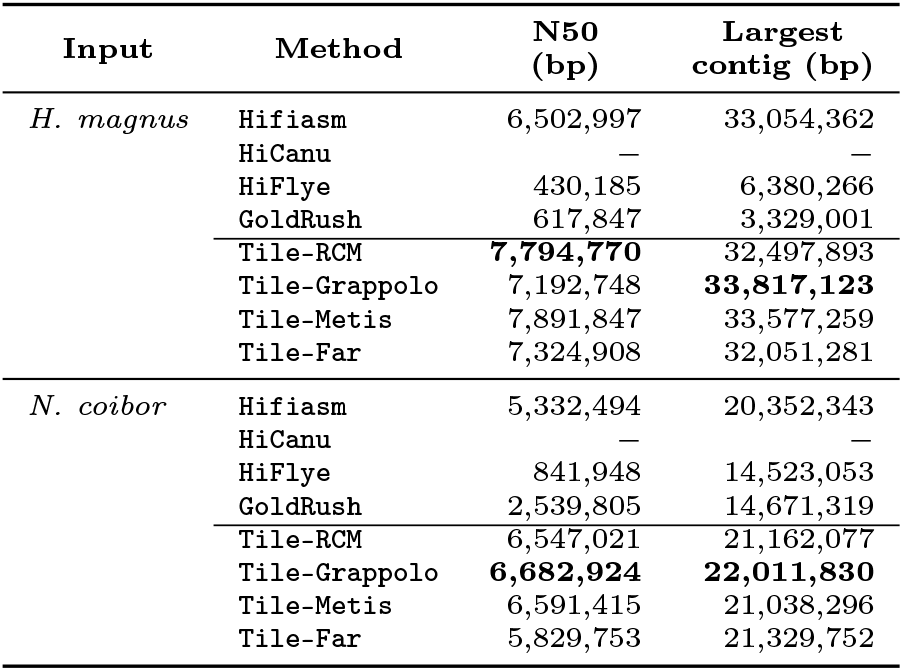
Real-world long read inputs analysis: Quality comparison of the output contigs generated by the different tools on two real-world inputs. Symbol − indicates that the corresponding runs required more than 252GB of memory, which was the system maximum memory. Boldface values show the best results for the given input.

## 4 Conclusion

In this paper, we revisited the long read assembly problem through the lens of read reordering. We presented `Tile-X`, a suite of algorithms that use various ordering schemes, to achieve both performance and qualitative gains. We also proposed a new variant of ordering that uses sparsification. Our findings indicate that (a) ordering helps reduce the computational burden of assembly for state-of-the-art assemblers; and (b) sparsified ordering delivers significant performance gains while preserving quality. Our current work does not harness the full potential of ordering yet—i.e., assembly performance can be improved if we use the pairwise ordering information to output the assembly without using a third party assembler— a direction that is of immediate future interest. Other future directions include: a) introducing a way to control sparsification rate and thereby associated trade-offs; b) providing qualitative guarantees with ordering for repetitive regions; and c) applying `Tile-X` on long read data sets obtained through a range of sequencing technologies.

## Acknowledgements

This research was supported in parts by NSF grants CCF 1919122 and CCF 2316160.

