## Supplementary Materials for "Tile-X: A vertex reordering approach for scalable long read assembly [Proceedings]"

### Supplementary Material

#### S1 Run-time breakdown

Figure S1 illustrates the execution time distribution across different phases of `Tile-X` for 4 inputs (2 simulated and 2 real-world), comparing the performance of `Tile-Grappolo` and `Tile-Far` schemes. The Figure S1(a) focuses specifically on the time spent in batching and subsequent steps, as mapping and graph construction account for 50-60% of the total execution time, overshadowing the time-saving benefits of sparsification in the overall performance comparison shown in Table 3.

Here, `Tile-Far` demonstrates a substantial reduction in execution time for the batched assembly and `iterative contig elongation` phase compared to `Tile-Grappolo`. This reduction directly correlates with the significant sparsification achieved by `Tile-Far`, as shown in Figure S1(b). For `Tile-Grappolo`, all long reads are retained for batching, whereas `Tile-Far` utilizes only 50-60% of the total reads, effectively reducing the computational burden. This significant reduction directly impacts the computational cost in subsequent phases.

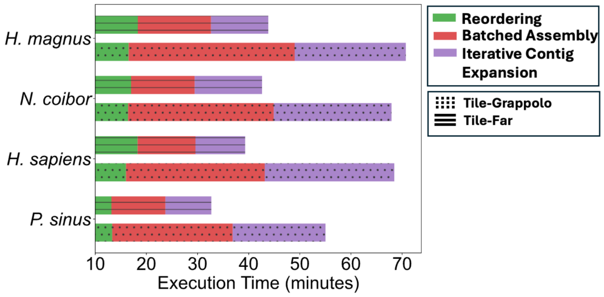


(a)

| Input | # reads in input graph | # of reads used in batched assembly | |
|---|---|---|---|
| | | Tile-Grappolo | Tile-Far |
| *P. sinus* | 2,323,801 | 2,323,801 | 1,580,185 |
| *H sapiens* | 3,083,048 | 3,083,048 | 2,065,642 |
| *H.magnus* | 1,919,461 | 1,919,461 | 1,190,064 |
| *N. coibor* | 2,436,589 | 2,436,589 | 1,453,707 |

(b)

Figure S1: (a) Execution time distribution across reordering, batched assembly, and `iterative contig elongation` comparing `Tile-Grappolo` and `Tile-Far` for 4 inputs (2 simulated and 2 real-world). Both the simulated inputs (*P. sinus* and *H. sapiens* are of 10x coverage); (b) Table summarizing the number of reads in the input graph and number of reads used in the batched assembly for both `Tile-Grappolo` and `Tile-Far`.

| Input | Method | Time Taken (in mins) | Peak Memory (in GB) |
|---|---|---|---|
| *D. busckii* | Hifiasm | 8.55 | 20.34 |
| | HiCanu | 15.01 | 28.03 |
| | HiFlye | 22.22 | 46.31 |
| | GoldRush | 36.03 | 51.72 |
| | Tile-RCM | 4.04 | 20.27 |
| | Tile-Grappolo | 3.51 | 22.29 |
| | Tile-Metis | 4.02 | 19.64 |
| | Tile-Far | **3.49** | **12.55** |
| *C. septempunctata* | Hifiasm | 34.3 | 28.07 |
| | HiCanu | 50.06 | 69.21 |
| | HiFlye | 48 | 73.07 |
| | GoldRush | 143.13 | 79.57 |
| | Tile-RCM | 12.42 | 18.31 |
| | Tile-Grappolo | 10.33 | 20.02 |
| | Tile-Metis | 10.33 | 20.32 |
| | Tile-Far | **9.55** | **14.41** |
| *B. splendens* | Hifiasm | 32.4 | 37.54 |
| | HiCanu | 20.18 | 64.10 |
| | HiFlye | 89.48 | 80.29 |
| | GoldRush | 161.01 | 72.33 |
| | Tile-RCM | 18.43 | 22.29 |
| | Tile-Grappolo | 15.29 | 24.36 |
| | Tile-Metis | 15.16 | 24.01 |
| | Tile-Far | **13.00** | **14.70** |
| *H. aestivaria* | Hifiasm | 81.98 | 48.07 |
| | HiCanu | 94.05 | 79.23 |
| | HiFlye | 143.38 | 83.55 |
| | GoldRush | 159.35 | 76.28 |
| | Tile-RCM | 57.33 | 29.19 |
| | Tile-Grappolo | 53.09 | 27.3 |
| | Tile-Metis | 54.41 | 29.5 |
| | Tile-Far | **35.33** | **18.25** |
| *P. sinus* | Hifiasm | 130.54 | 54.62 |
| | HiCanu | 166.25 | 231.03 |
| | HiFlye | 313.29 | 107.40 |
| | GoldRush | 210.02 | 92.27 |
| | Tile-RCM | 107.05 | 27.08 |
| | Tile-Grappolo | 103.54 | 29.09 |
| | Tile-Metis | 101.39 | 29.11 |
| | Tile-Far | **81.21** | **20.25** |
| *H. sapiens* | Hifiasm | 364.94 | 71.02 |
| | HiCanu | * | * |
| | HiFlye | - | - |
| | GoldRush | 409.12 | 108.07 |
| | Tile-RCM | 228.51 | 41.29 |
| | Tile-Grappolo | 220.34 | 42.21 |
| | Tile-Metis | 222.13 | 41.26 |
| | Tile-Far | **191.31** | **21.45** |

Table S1: Performance comparison of the output contigs generated by the different tools on the different inputs. Symbol $*$ indicates that the corresponding runs did not complete within 6 hours; symbol $-$ indicates that the corresponding runs required more than 252GB of memory which was the system maximum memory. Bold face values show the best results for any input.
